# Effect of Potassium Nitrate and Calcium Chloride priming on germination and seedling growth of Honesty (*Lunaria annua* L.)

**DOI:** 10.1101/2022.12.02.518881

**Authors:** Reyaz Ahmad Bhat, F U Khan, Z. A. Bhat, F. A. Khan, Javaid Ahmad Wani, Abu Manzar, I. T Nazki, Neelofar, Qazi Altaf, Nasir Hamid Masoodi

**Author notes:** Corresponding author’s email id.

## Abstract

Seed priming improves seed performance under environmental conditions. The study was designed to evaluate the effect of different priming treatments on the germination behaviour of Honesty (Lunaria annua L.). The experiment was conducted under a complete randomized design (CRD) with three replications. Seed priming was done with different concentrations of calcium chloride (CaCl_2_), potassium nitrate (KNO3) and hydro priming. All the treatments except T_1_ (Control) had a significant effect on seedling establishment and seedling vigour, Results show that maximum invigoration was observed in seeds primed at 1% KNO3 while minimum invigoration was observed in T_1_ (Control). It was concluded that the germination percentage could be increased by using lower concentrations of KNO3 and CaCl_2_.

## INTRODUCTION

Honesty (*Lunaria annua* L.), that is also known as silver dollar, satin flower, money plant, and moonwort, belongs to family Brassicaceae. It’s a tall biennial ornamental with hairy stems that’s native to southeastern Europe and western Asia. It has unstalked, coarsely serrated leaves with huge scentless flowers of 25-30 mm in diameter that bloom from April to June (Fitter *et al*., 1980) if the Vernalization conditions have been met throughout the previous fall and winter (Pierik,1967). a biennial crop that starts growing in the spring and stays vegetative for the first year. Lunaria is a spring flowering plant with toothed leaves and odorless flowers that grows without petioles (April to June). This plant, in addition to its attractiveness, produces lovely violet and white flowers that may be used as decorative plants at home for several years after the fruits (coins) have reached full maturity and the fruit inflorescence has been harvested. The skeletal appearance of the Lunaria, which is most stunning in its dried condition, is most eye-catching in the winter garden. Lunaria gets its common and botanical names from the oval seed pods and the gleaming silique that remains once the pod splits to release the seed.

Seed germination is considered as critical stage in the whole life of a plant (Yang et al., 2008). Poor seed germination and uneven seedling emergence results into low quality products which leads to financial losses to growers. Several techniques and strategies have been developed to improve seed germination and uniform seedling growth. Seed priming is a presoaking seed treatment to improve seed germination and uniform seedling emergence in a wide range of crops (McDonald, 2000). In seed priming, seeds are partially hydrated so that it activates metabolic activities but prevents radicle protrusion. After seed priming, seeds are re-dried to initial moisture content prior to sowing

Osmotic adjustment or priming of seeds prior to sowing is known as an efficient way to increase germination and emergence rate in some species with for example small embryo or species with stepwise seed development (Sivritepe, 2000). The beneficial effects of priming have also been demonstrated for many field crops (Mehmet *et al*., 2006). Seed priming has been successfully demonstrated to improve germination and emergence in seeds of many crops, particularly seeds of vegetables and small seeded grasses (Arif *et al*., 2007). The purpose of these treatments is to shorten the emergence and to protect seed from biotic and abiotic factor during critical phase of seedling establishment so as to synchronize emergence, which lead to uniform stand and improved yield. These priming treatments which enhance seed germination include hydro-priming (Basra *et al*., 2003; Afzal *et al*., 2004), osmopriming and hormonal priming (Afzal *et al*., 2006). It was indicated that primed seed emerged one to three days earlier than non primed seeds and quickly became apparent. Similarly, improvement in germination, reduction in germination time and enhanced emergence in hydro-primed seeds were reported by Harris *et al*. (2001). Priming is the enhancement of physiological and biochemical events in seeds during suspension of germination by low osmotic potential and negligible matric-potential of the imbibing medium. Salts or non penetrating organic solutes in liquid medium (osmo-conditioning) or matrix solutions (matri-conditioning) are used to establish equilibrium water potential between seed and the osmotic medium needed for conditioning (Khan, 1993). The present study was planned to evaluate the effects of KNO3 and CaCl_2_ on seed germination of Lunaria annua L.

## 2. MATERIALS AND METHODS

Experiment was conducted in laboratory of division of floriculture and landscape architecture faculty of Horticulture, Sher-e-Kashmir university of Agriculture Sciences Srinagar Kashmir, during the year 2021 To improve the rate of germination and reduce the time required for germination, seed priming was done with different concentration of calcium chloride (CaCl_2_) potassium nitrate (KNO3) and hydro-priming to determine its effect on seed germination (Table 1 & 2). For priming, seeds were soaked in distilled water and different concentration of potassium nitrate solution for 48 h. After treatment, the seeds were placed on wet filter paper treated with fungicide Dithane M-45 @ 1 g/L in growth room. The seeds were kept in growth chamber at 24 ± 2°C. The seeds were watered at three days interval and kept under close observation. Treatments were designed with three replications and 10 seeds per replication in complete randomized design (CRD). Experimental data was analyzed statistically adopting the techniques of analysis of variance (ANOVA) for Completely Randomized Design (CRD). The level of significance of the treatment mean square at 5% probability was tested against F calculate value for Completely Randomized Design (CRD). The data were recorded on days to 50 % germination, germination percentage, seedling fresh weight, seedling dry weight, root length, shoot length, seedling vigour index and root/shoot ratio.

**Table 1.** Effect of different priming treatments on days to 50% germination, germination percentage, seedling fresh weight & seedling dry weight of Lunaria annua L.

| **Treatment combination** | **Days to 50% germination** | **Germination percentage** | **Seedling Fresh weight** | **Seedling dry weight** |
| --- | --- | --- | --- | --- |
| **T1:** Control | 14.6 | 40.0 | 102.8 | 19.1 |
| **T2:** CaCl<sub>2</sub> (1 %) | 12.9 | 56.2 | 108.6 | 21.2 |
| **T3:** CaCl<sub>2</sub> (2 %) | 11.7 | 58.3 | 110.0 | 22.9 |
| **T4:** CaCl<sub>2</sub> (3 %) | 11.6 | 61.1 | 114.2 | 24.6 |
| **T5:** CaCl<sub>2</sub> (4 %) | 12.8 | 54.0 | 109.0 | 22.7 |
| **T6:** KNO<sub>3</sub> (1 %) | **10.0** | **72.4** | **116.5** | **25.7** |
| **T7:** KNO<sub>3</sub> (2 %) | 11.0 | 69.3 | 114.6 | 25.6 |
| **T8:** KNO<sub>3</sub> (3 %) | 11.3 | 68.1 | 111.9 | 24.3 |
| **T9:** KNO<sub>3</sub> (4 %) | 12.3 | 67.3 | 112.3 | 24.1 |
| **C.D.** | **1.39** | **4.09** | **7.48** | **2.92** |
| **SE(m)** | **0.46** | **1.36** | **2.50** | **0.97** |

**Table 2.** Effect of seed priming treatments on Seedling dry weight, root length, shoot length, Seedling vigour index and root/shoot ratio.

| **Treatment combination** | **Root length (cm)** | **Shoot length (cm)** | **Seedling vigour index** | **Root/shoot ratio** |
| --- | --- | --- | --- | --- |
| **T1:** Control | 3.5 | 3.0 | 332.4 | 0.33 |
| **T2:** CaCl<sub>2</sub> (1 %) | 5.4 | 3.9 | 419.1 | 1.02 |
| **T3:** CaCl<sub>2</sub> (2 %) | 5.6 | 4.0 | 516.6 | 0.59 |
| **T4:** CaCl<sub>2</sub> (3 %) | 6.1 | 4.1 | 550.4 | 0.73 |
| **T5:** CaCl<sub>2</sub> (4 %) | 5.7 | 4.3 | 510.0 | 0.91 |
| **T6:** KNO<sub>3</sub> (1 %) | **6.9** | **7.1** | **670.0** | **1.09** |
| **T7:** KNO<sub>3</sub> (2 %) | 6.4 | 6.3 | 636.6 | 1.01 |
| **T8:** KNO<sub>3</sub> (3 %) | 5.8 | 4.7 | 590.3 | 1.02 |
| **T9:** KNO<sub>3</sub> (4 %) | 5.7 | 4.4 | 603.3 | 0.82 |
| **C.D** | **1.19** | **0.89** | **128.3** | **0.36** |
| **SE(m)** | **0.40** | **0.29** | **42.86** | **0.12** |

## 3. RESULTS

### 3.1 Seedling Establishment

Seedling establishment of *Lunaria annua* L. includes number of days required by seeds to 50% germination (%) and Germination percentage. The results of the present study regarding days to 50% germination & germination percentage presented in the Table 1 shows statistically significant differences among different treatments but the seed priming with KNO3 improved the stand establishment of Honesty (*Lunaria annua* L.) grown in growth chamber. Honesty (*Lunaria annua* L.) seeds primed with 1% KNO3 had maximum emergence rate (76.3 %), so they were showing better performances when compared to the other treatments. Minimum germination (40%) was observed in non treated I.e. T_1_ (control) seeds of Honesty (*Lunaria annua* L.), while, in the case of days to 50% germination, the maximum number of days (14.6) was observed in non treated seeds of *Lunaria annua* L. The seeds treated with 1% KNO3 germinated earlier than all other treatments.

### 3.2 Seedling Vigor

Statistical analysis of data about seedling vigor revealed that all priming treatments significantly improved the seedling shoot length in Honesty (*Lunaria annua* L.), whereas the maximum seedling root and shoot length was achieved in Lunaria seed primed with 1% (6.9 & 7.1 cm), followed by 2% KNO3 solution (6.4 & 6.3 cm), whereas the lowest values for seedling root length and shoot length was observed in control (3.5 & 3 cm)., plants raised from seeds treated with 1% KNO3 showed higher values for seedling fresh weight (116.5 mg) and dry weight (25.7 mg) respectively as compared to other treatments (Table 2).

**Figure1.**
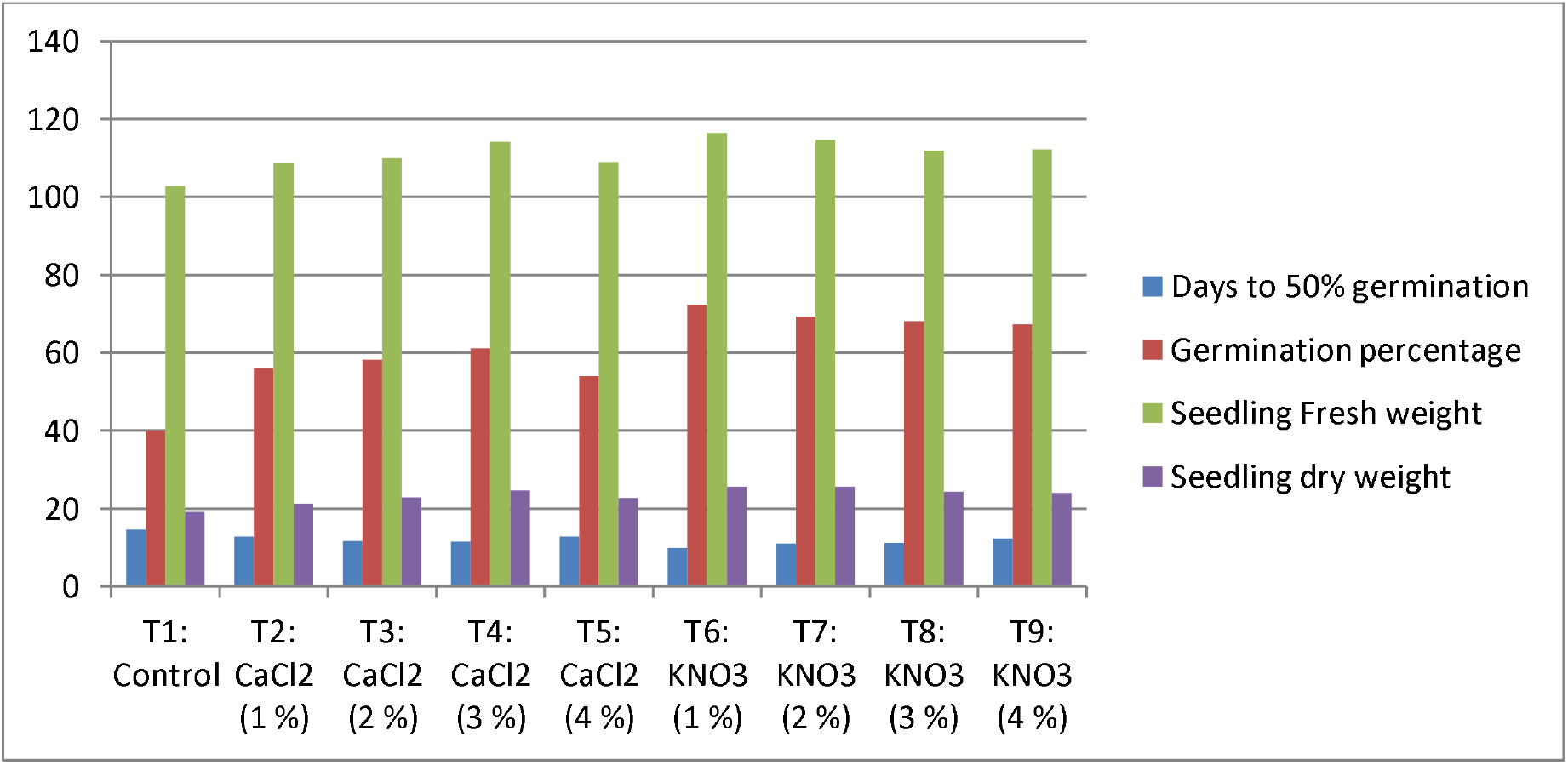
Graphical representation of effect of different priming treatments on various seedling establishment parameters.

**Figure2.**
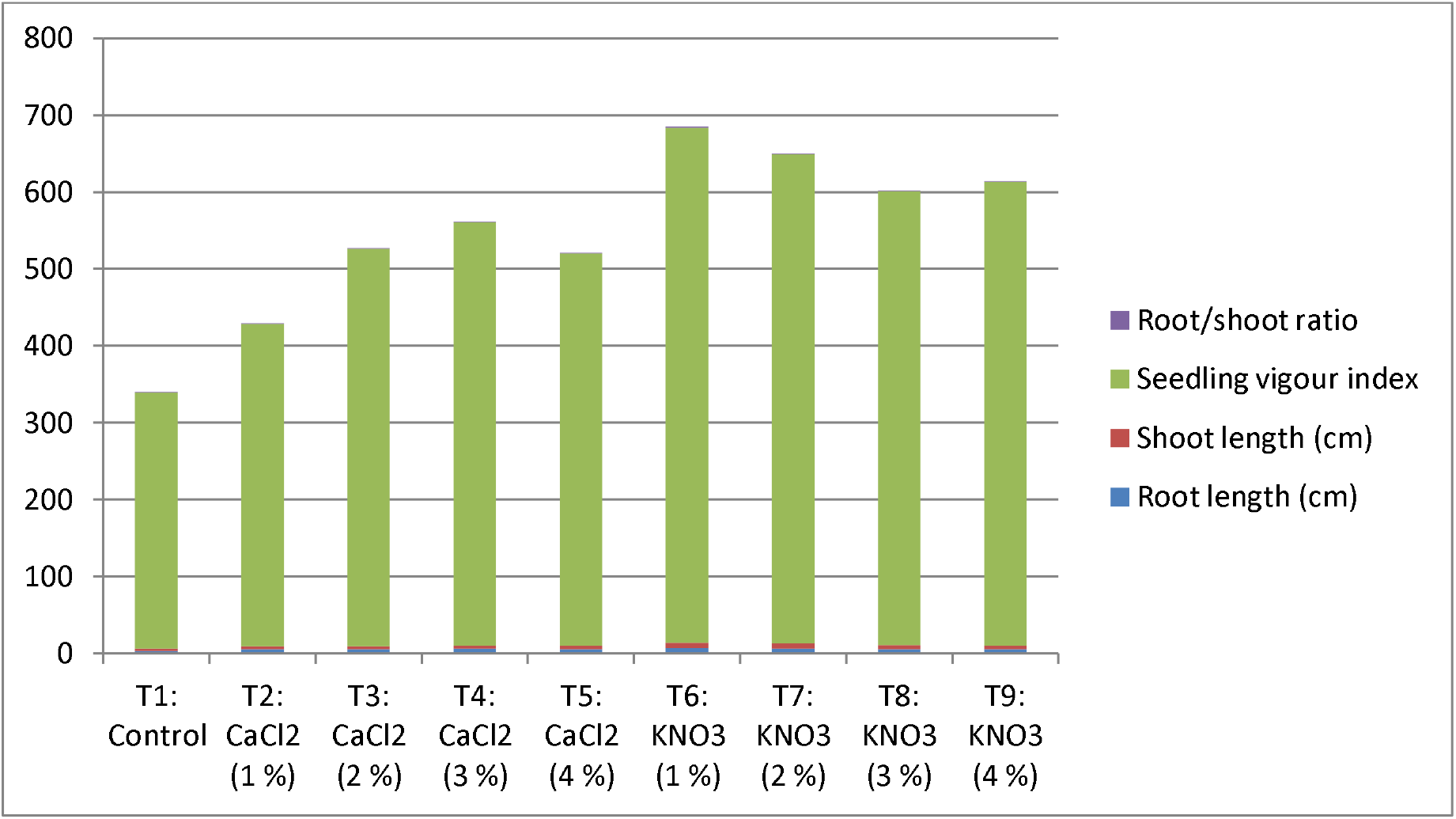
Graphical representation of effect of different priming treatments on various seedling vigor parameters.

## 4. DISCUSSION

### 4.1 Seedling establishment

Honesty (*Lunaria annua* L.) seed priming with different concentrations of KNO3 and CaCl_2_ affected the emergence of seedling and the speed of seed germination. Major events in other literature on priming includes metabolic changes, such as repair of DNA and increases in the biosynthesis of RNA (Bray 2017), and enhancement in the respiration process of seed (Singh *et al*., 2013). This indicates that the time of seed imbibition is very important for seed priming. For the study of seed priming of Honesty (*Lunaria annua* L.) with different levels of KNO3 and CaCl_2_, it is important to know about the emergence percentage and emergence time.

The data shown in Table 1 indicated that priming of Honesty (*Lunaria annua* L.) seeds with 1% KNO3 was better than other treatments in terms of days to 50% germination and germination percentage. The mechanism of action of KNO_3_ on the improvement of seed germination and/or early growth is far from being completely understood. The complexity about nitrate effects on seed germination and early seedling growth could be due to its dual role as a nutrient and a signaling molecule (Duermeyer *et al*., 2018). Nitrate stimulation of seed germination is often associated with plant species whose seeds require light for germination (Hilhorst *et al*., 1986: Footitt *et al*., 2013). Our study is in correspondence with another study that revealed that the emergence percentage of wheat seeds was decreased when they were primed with >1% KNO3 (Shafiei *et al*., 2012). This indicates that KNO3 concentration above a certain threshold may not be appropriate to boost seed germination. Seed priming with 1% KNO3 was found useful in terms of emergence percentage in sorghum (Shehzad *et al*., 2012) and rice (Ruttanaruangboworn 2017). Besides, soybean seed primed with 1% KNO3 for 1 day enhanced the emergence percentage as compared to non-treated seeds, both in laboratory and field experiments (Mohammadi 2018).

### 4.2 Seedling Vigor

Seedling vigor is the combined result of the emerged seeds under a wide range of biotic and abiotic stresses. Seedling vigor is not a single measurable entity, but it is a sum of many growth parameters, such as seedling length, seedling fresh weight, and seedling dry weight (ISTA 2015). Maximum vigor was observed when seed priming with 1% KNO3 was done. Our study is in line with another study in which seedling vigor of wheat was improved by priming with KNO3 (Shafiei *et al*., 2012). Similar results were found in corn when the priming of seed was done with 1% KNO3 (Hadinezhad *et al*., 2013). Our findings are similar to other studies, in which the shoot length of watermelon and tomato were increased by the seed priming with KNO3 (Demir *et al*., 1999 & Mirabi 2012). Seed priming with 1% KNO3 improved the vegetative growth of watermelon (Oliveira *et al*., 2019) and tomato (Vaktabhai *et al*., 2017), respectively, under salt stress. Seed priming with KNO3 can cause a significant increase in seedling vigor of the wheat crop as compared to hydro-priming or dry broadcasting (Basra *et al*., 2006)

## 5. Conclusion

The performance of honesty is diminished by the poor quality of seed. Therefore, the present study was conducted to improve the quality of honesty seed by priming with different concentrations of KNO3 & CaCl_2_. The results presented in this paper revealed that honesty seeds primed with 1% KNO3 proved to be successful for improving seedling establishment and vigor. The present study provides the direction towards further molecular investigation related to the seed priming of honesty.

**Fig.3.**
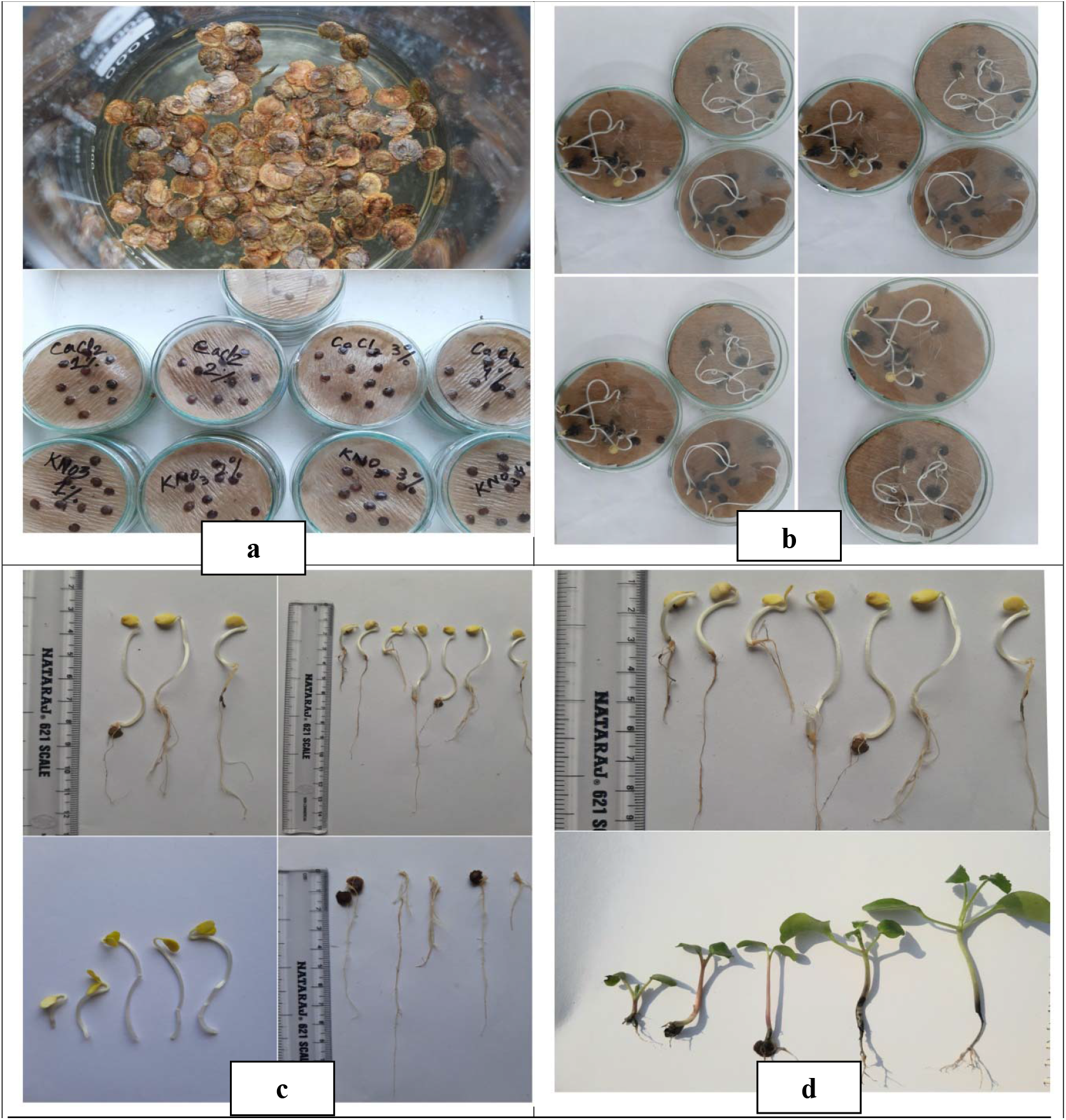
**a)** Primed honesty seed in Petri-plates **b)** 50% seed germination **c, d)** measuring different seedling parameters

